## Supplementary Figures for "Paired yeast one-hybrid assays to detect DNA-binding cooperativity and antagonism across transcription factors"

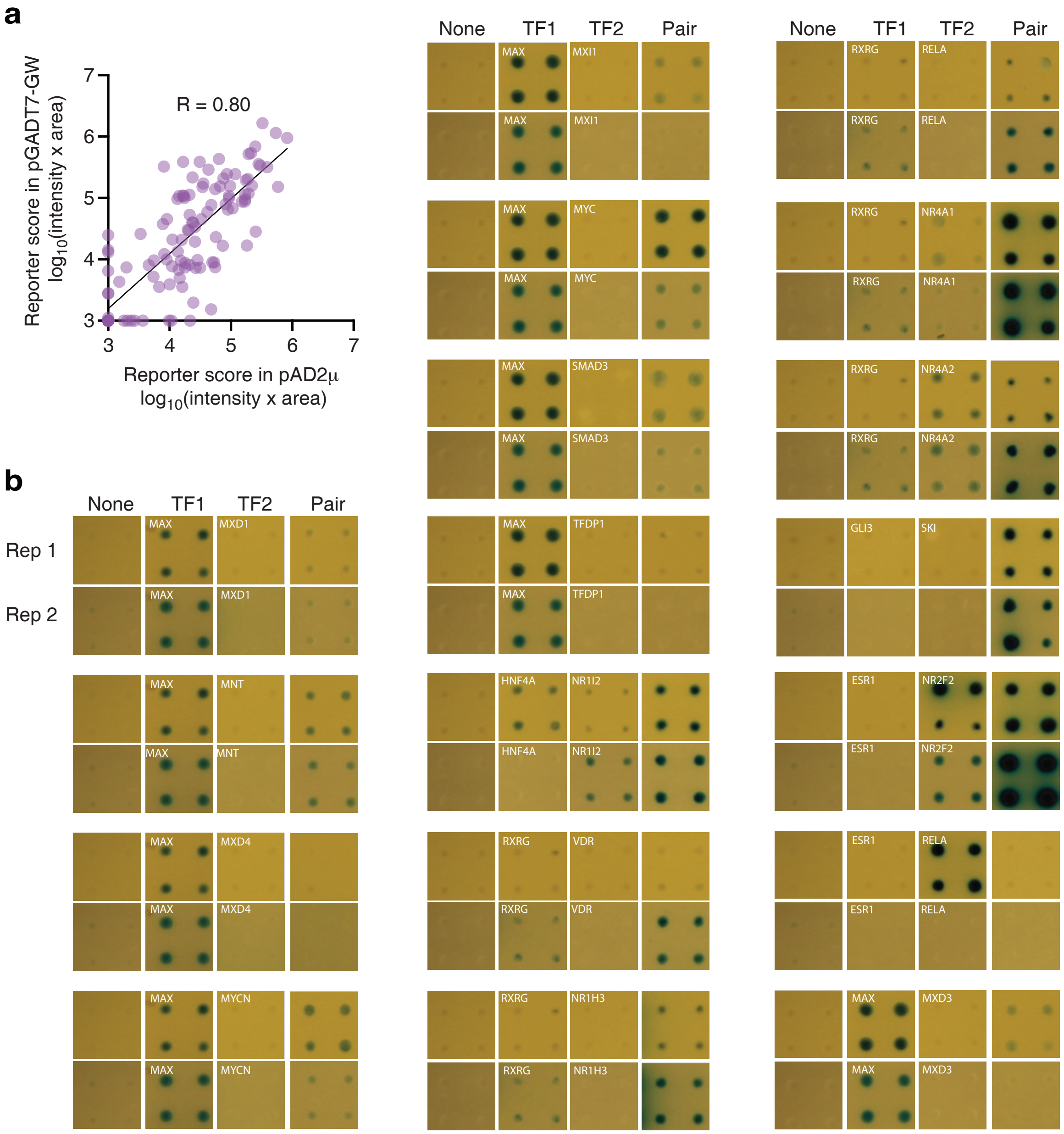


**Supplementary Figure 1. Reproducibility of pY1H assays.** (**a**) Correlation between reporter scores (intensity x area) for single-TFs cloned into the pGADT7-GW and the pAD2μ vectors and transformed into yeast. (**b**) Cooperative and antagonistic interactions from two pY1H screens using rthe 2-AD design conducted on different days for the CCL15 promoter.

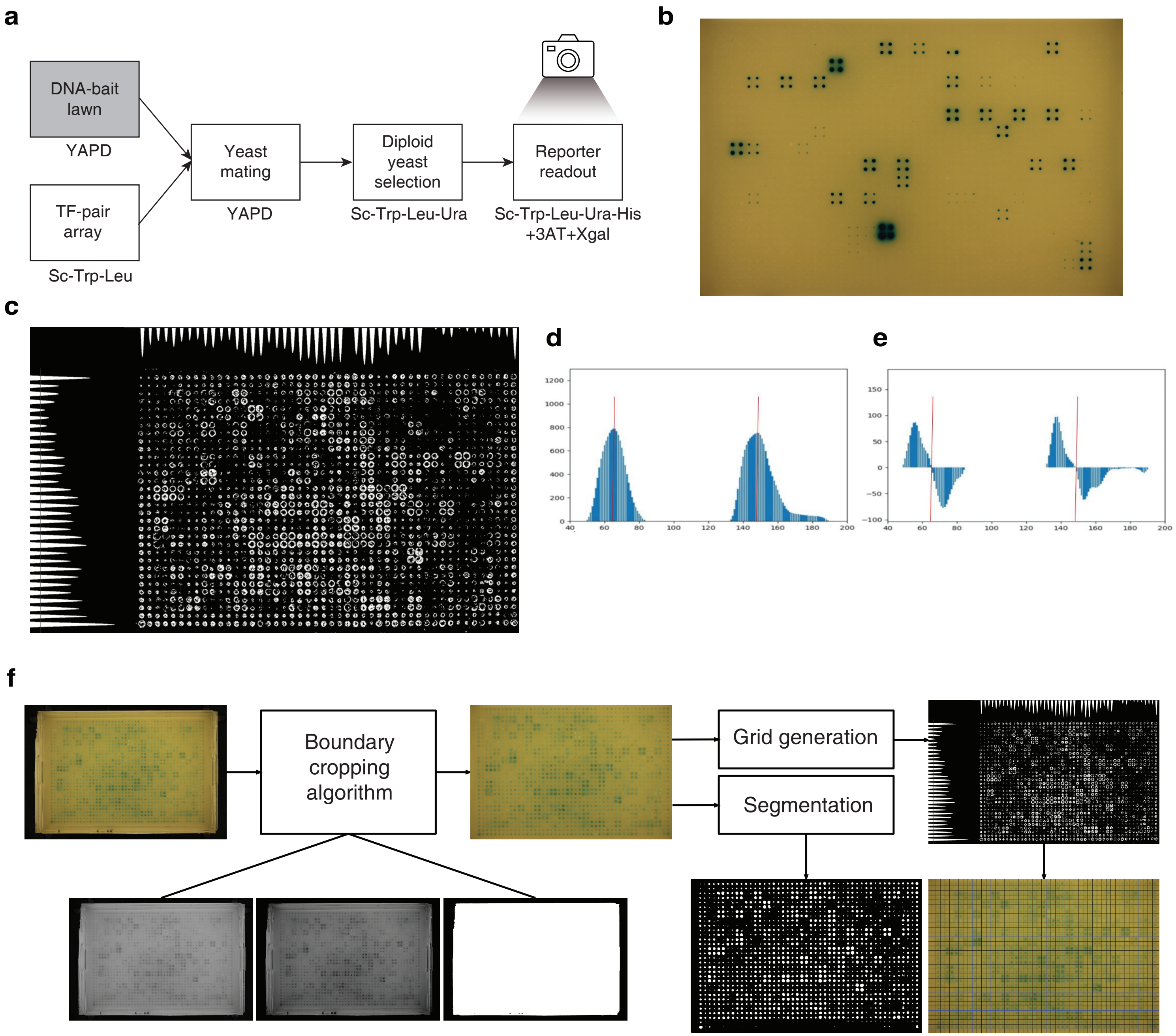


**Supplementary Figure 2. pY1H experimental and analysis pipeline.** (**a**) Experimental pipeline for pY1H assays. A lawn of the yeast DNA-bait is mated with the TF-pair array in a YAPD plate for 1 day. Diploid yeast are selected in Sc-Trp-Leu-Ura plates for 2 days and then transfered to readout plates containing 3AT and X-gal. Pictures are taken at days 2, 3, 4 and 7. (**b**) Example of a pY1H readout plate. (**c**) Vertical and horizontal histogram projection of a sub-optimal segmentation mask. Each peak corresponds to the center of the colony. (**d**) Plot of a 160-pixel region of histogram projection. (**e**) Plot of the first-order derivative of the region of the plot in d. Each “zero-crossing” event corresponds to the histogram's peak or the colony's center. (**f**) Pipeline of the image analysis process. First, the boundary cropping algorithm selects the region of interest. Then the grid generation algorithm performs histogram projection analysis to generate a grid based on the TF-pair array coordinates. Finally, the segmentation model generates a segmentation map for the colonies and determine colony area and intensity of blue color.

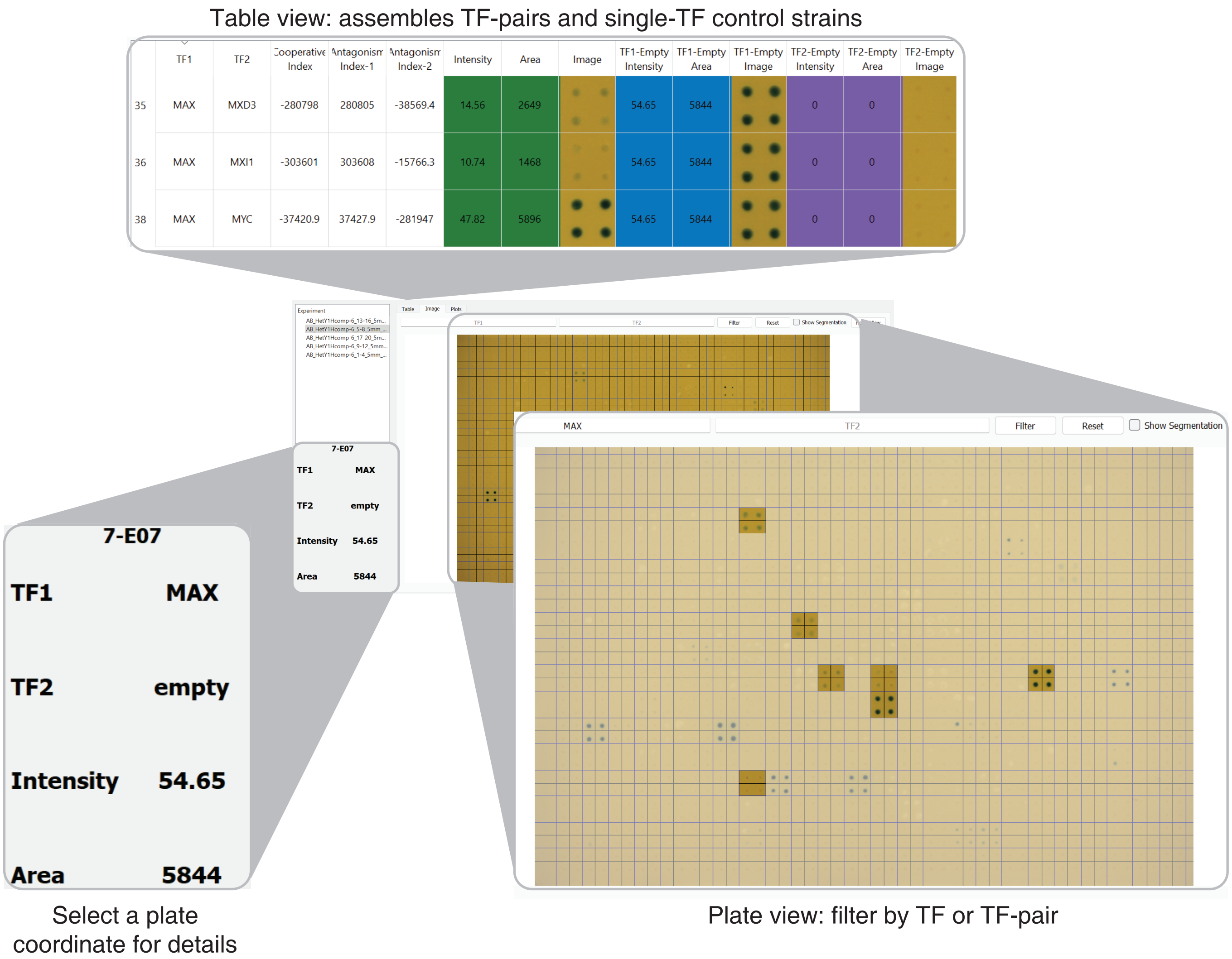


**Supplementary Figure 3. Features of DISHA.** The Plate view displays plate images derived from pY1H assays and allows to filter by singe-TF or TF-pair. Users can also select an array coordinate to identify the TF-pair, reporter intensity measured as blue color, and area of colonies. The Table view displays images of TF-pair colonies alongside single-TF colonies allowing side-by-side comparisons, together with reporter intensity and colony area. In addition, the table presents the cooperative and antagonistic indices calculated for each TF-pair.

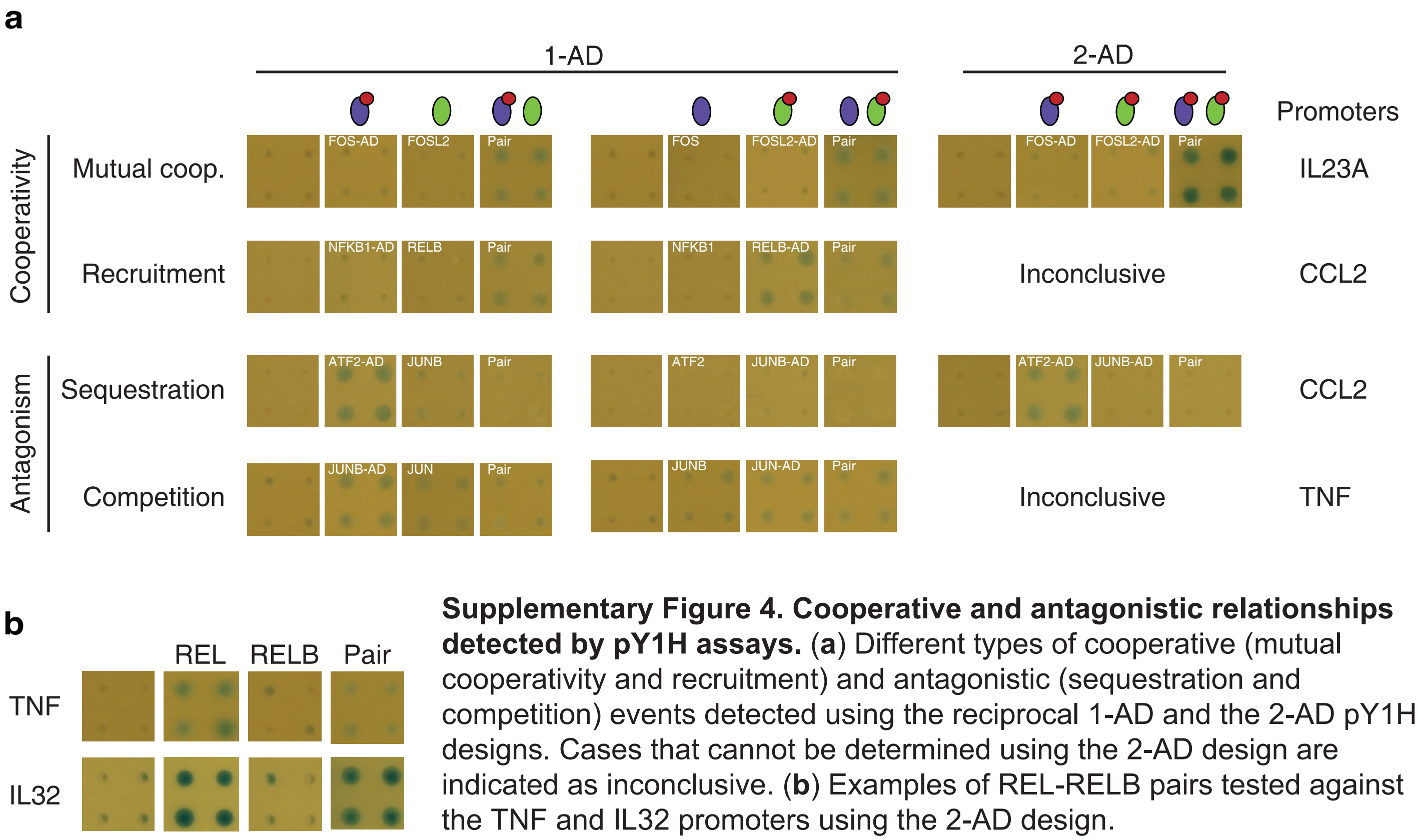


**Supplementary Figure 4. Cooperative and antagonistic relationships detected by pY1H assays.** (**a**) Different types of cooperative (mutual cooperativity and recruitment) and antagonistic (sequestration and competition) events detected using the reciprocal 1-AD and the 2-AD pY1H designs. Cases that cannot be determined using the 2-AD design are indicated as inconclusive. (**b**) Examples of REL-RELB pairs tested against the TNF and IL32 promoters using the 2-AD design.

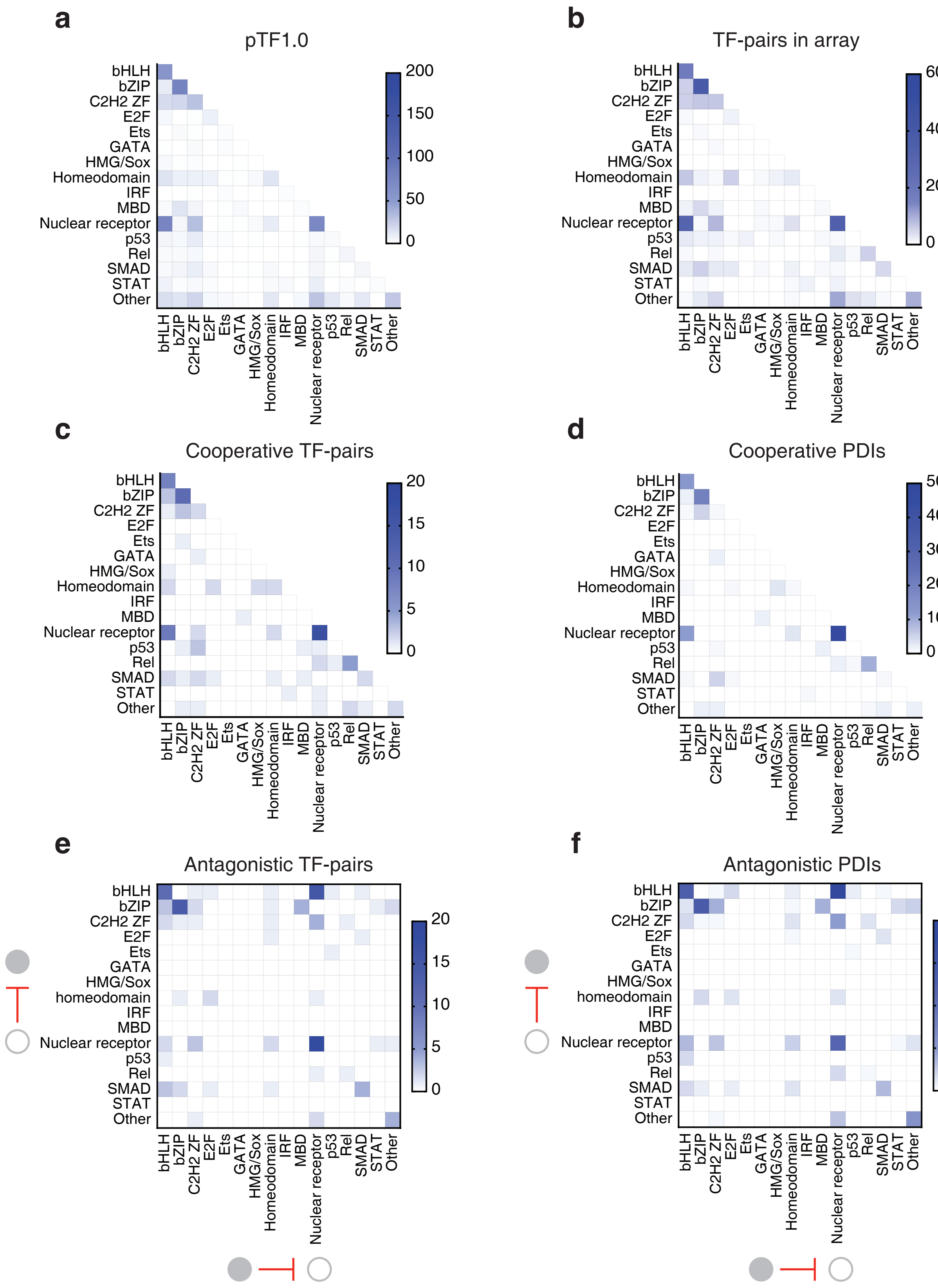


**Supplementary Figure 5. TF family representation.** (**a, b**) Number of TF-pairs for each TF family combination for the 868 TF-pairs in pTF1.0 (**a**) and the 297 sequence-confirmed TF-pairs in the array (**b**) (as in Fig. 3b,c). (**c, d**) Number of cooperative TF-pairs (**c**) and PDIs (**d**) identified in a pY1H assay against 18 cytokine promoters. (**e, f**) Number of antagonistic TF-pairs (**e**) and PDIs (**f**) identified in a pY1H assay against 18 cytokine promoters. Columns indicate the family of the antagonistic TF and rows indicate the family of the antagonized TF.

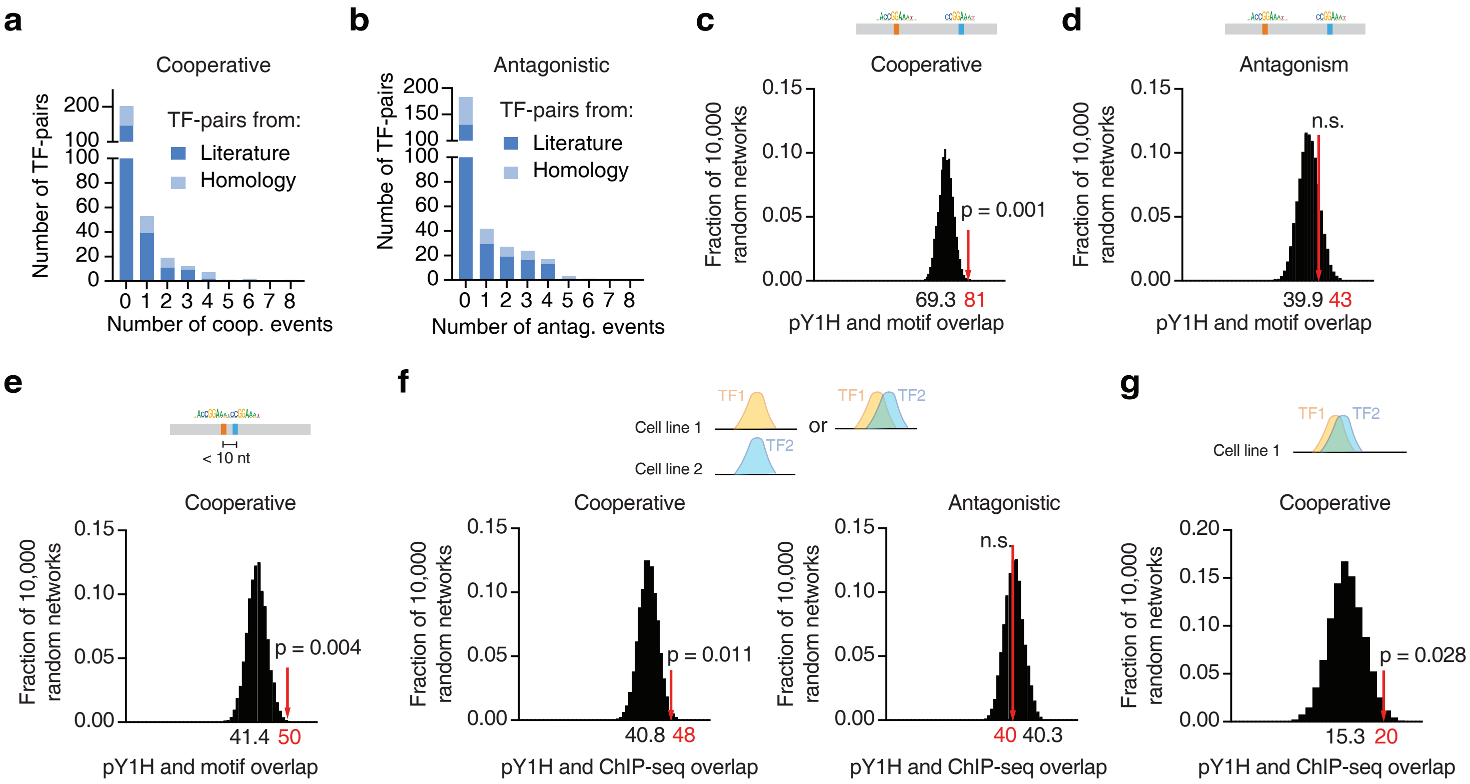


**Supplementary Figure 6. Cooperative and antagonistic events.** (**a**, **b**) Number of TF-pairs involved in different number of cooperative (**a**) or antagonistic events for TF-pairs derived from the literature or predicted based on homology (**b**). (**c, d**) Overlap between pY1H cooperative (**c**) and antagonistic (**d**) events with the presence of motifs for both single-TFs in the corresponding promoter sequences. The histograms show the distribution of the overlap with each randomized pY1H network, where the pY1H network was randomized 10,000 times by edge-switching (each edge is a TF-pair and a cytokine promoter). The numbers under the histograms indicate the average overlap in 10,000 randomized networks, while the red arrows indicate the observed overlap with the actual pY1H-derived network. Statistical significance was calculated from Z-score values assuming normal distribution for overlap with the randomized networks. (**e**) Overlap between pY1H cooperative events with the presence of motifs for both single-TFs within 10 bp in the corresponding promoter sequences. (**f, g**) Overlap between pY1H cooperative or antagonistic events and the presence of ChIP-seq peaks from GTRD for both single-TFs in the corresponding promoter sequences. ChIP-seq peak summits were required to be within 50 bp irrespective of the cell line (**f**) or in the same cell line (**g**).

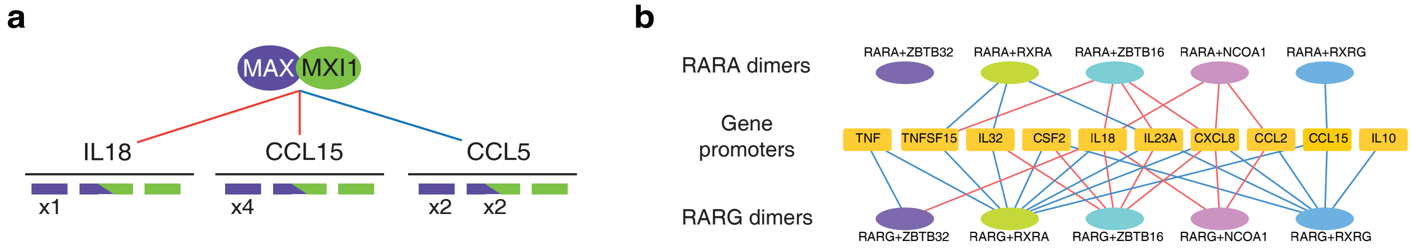


**Supplementary Figure 7. DNA binding specificity for T-pairs.** (**a**) Number of distinct and overlapping motifs for MAX and MXI1 in the indicated cytokine gene promoters. (**b**) Cooperative and antagonistic events for TF-pairs involving paralogs RARA and RARG.
